# Integrating AlphaFold2 Models and Clinical Data to Improve the Assessment of Short Linear Motifs (SLiMs) and Their Variants’ Pathogenicity

**DOI:** 10.1101/2025.01.27.634988

**Authors:** Franco G. Brunello, Lorenzo Erra, Juan Nicola, Marcelo A. Marti

## Abstract

Short Linear Motifs (SLiMs) are protein functionally relevant regions that mediate reversible protein-protein interactions. Variants that disrupt SLiMs can lead to numerous Mendelian diseases. Although various bioinformatic tools have been developed to identify SLiMs, most suffer from low specificity. In our previous work, we demonstrated that integrating sequence variant information with structural analysis can enhance the prediction of true functional SLiMs while simultaneously generating tolerance matrices that indicate whether each of the 19 possible single amino acid substitutions (SASs) is tolerated. However, the scarcity of representative crystallographic structures of SLiM-receptor complexes posed a significant limitation. In this study, we demonstrate that these interactions can be modeled using AlphaFold2 (AF2) to generate high-quality structures that serve as input for our MotSASi method. These AF2-derived structures show robust performance, both in reproducing known structures deposited in the Protein Data Bank (PDB) and in reflecting the deleterious effects of known sequence variants. This updated version of MotSASi expands the repertoire of high-confidence predicted SLiMs and provides a comprehensive catalog of variants located within SLiMs, along with their respective deleteriousness assessments. When compared to AlphaMissense, MotSASi demonstrates superior performance in predicting variant deleteriousness. By contributing to the accurate identification and interpretation of variants, this work aligns with ACMG/AMP standards and aims to improve diagnostic rates in clinical genomics.

**Author Summary:** Proteins interact with each other in highly specific ways to carry out vital biological functions. Short Linear Motifs (SLiMs) are small regions within proteins that mediate many of these reversible interactions. Changes in SLiMs can disrupt these interactions and lead to severe genetic disorders. Identifying SLiMs accurately has been a longstanding challenge, as many computational tools suffer from low specificity. Previously, we developed a method, MotSASi, that combines sequence variation data and structural analysis to improve SLiM prediction and assess the impact of single amino acid substitutions (SASs). However, the lack of available structural data limited its application. In this study, we demonstrate that structures generated using AlphaFold2 (AF2) can overcome this limitation. These high-quality AF2 models reliably reproduce known structures and capture the harmful effects of sequence variations. By integrating AF2 models, the updated MotSASi method identifies more high-confidence SLiMs and provides detailed assessments of the variants within them. MotSASi outperforms existing tools, such as AlphaMissense, in predicting the impact of genetic variants, offering insights aligned with clinical standards. This advancement can aid in understanding disease mechanisms and improving genetic diagnostics in clinical genomics.

## Introduction

Short Linear Motifs (SLiMs) are short stretches of amino acids found in protein sequences, often defined by regular expressions, and typically located in disordered regions of proteins or exposed flexible loops. These characteristics allow SLiMs to interact with their domain-binding partners reversibly and transiently, contributing to proper biological function of the protein host. SLiMs are crucial for normal cellular physiology, participating in various processes such as cell cycle, vesicle trafficking, cytoskeleton dynamics, innate immunity, and protein and RNA degradation systems, all of which play key roles in regulating cellular decision-making [1]. Given their significant biological roles, their identification and characterization are essential for understanding molecular cell biology. However, due to the difficulties and limitations of their experimental identification, only a small fraction of SLiMs have been deeply studied to date. More recently, there have also been some advances in medium- and high-throughput approaches for SLiM discovery [2, 3]. On the contrary, since most human (and many other) protein sequences are known, thousands of SLiMs have been bioinformatically predicted and await further scrutiny[4].

SLiM prediction and analysis are commonly approached using diverse bioinformatic tools[5–12], among which the universally recognized Eukaryotic Linear Motif (ELM) resource stands out. ELM is a comprehensive SLiM database and motif prediction tool [5]. In the latter role, it leverages information on sequence, structure, function, localization, evolutionary conservation, and interaction context to evaluate the confidence of motif-mediated protein complexes. Despite the mentioned advances, SLiM prediction remains impaired by very low specificity. This results in a high rate of false positives, which arise for several reasons, including the vastness of the search space (e.g., the whole human proteome) and the degenerate nature of many SLiM regular expressions, which makes their occurrence by chance, highly probable. Furthermore, the fact that many SLiM classes in the ELM database contain very few validated instances (i.e true positives) exacerbates the problem. In this context, downstream filtering strategies that improve SLiM prediction confidence are essential, which was the focus of our previous work, where we presented the Motif-occurring Single Amino acid Substitution information (MotSASI) method [13].

In this previous work, we incorporated two key approaches. First, we utilized crystallographic structures representing motif-domain interaction complexes (as derived from the PDB) to analyze the effects of single amino acid substitutions (SAS) within the motif, employing the FoldX software [14,15]. Specifically, we determined whether the motif was tolerant or intolerant to a given mutation based on the resulting change in the stability of the complex. Second, we evaluated the clinical significance (as annotated in ClinVar) and population allele frequency (as reported in gnomAD) of missense variants located within motif sequences [16,17]. In other words, we assessed whether the variants observed in the motif were classified as benign or pathogenic. The underlying premise was that non-functional SLiMs identified by chance through regular expression matching (false positives) exhibit a variation pattern—pathogenic (non-tolerant) versus benign (tolerant)—that differs significantly from that of functional SLiMs (true positives). Functional SLiMs display a SAS structural-clinical profile consistent with observed data. Through this approach, we were able to eliminate potential false positives and also increase the confidence in the identification of true predicted motifs.

The initial implementation of MotSASi yielded promising results, achieving enrichments ranging from 2- to 50-fold relative to the original true positive set (depending on the particularly analyzed motif), while also discarding a substantial number of false positives. However, the algorithm was constrained to classify candidates carrying known variants within motif classes with representative motif-receptor crystallographic structures, thereby limiting the analysis to only 10–20% of all motif matches in the proteome. Since the deployment of our first version, gnomAd and ClinVar databases have continued to expand, and the increasing availability of genomes and exomes deposited in gnomAD suggests significant potential for improved outcomes. More importantly, recent advances in machine learning approaches for protein structure prediction, such as AlphaFold2 as a key reference, have completely changed the protein structure landscape. These developments promise to bridge the gap that previously prevented the study of many more motif classes.

AlphaFold2 (AF2), the state-of-the-art AI system developed by Google DeepMind, computationally predicts protein structures with unprecedented accuracy and speed [18]. The system takes a query sequence as input and predicts first residue-residue contacts, and then the resulting structure by combining the construction of multiple sequence alignments. More recently, the user interface has been enhanced to support the incorporation of multiple simultaneous query chains, seamlessly aligning with our objective of modeling motif-domain interaction pairs [19].

In the present study, we leveraged AlphaFold’s structural prediction capabilities to generate models of all human SLiMs in complex with their domain-binding partners for motif classes that lacked representative crystallographic structures in the PDB. By integrating this structural data with population allele frequency information from gnomAD and clinically relevant variants from ClinVar, alongside a series of logical filters, we refined sequence-based predictions of functional SLiMs in the human proteome, resulting in a substantial number of high-confidence candidates, as well as many discarded potential false positives. Simultaneously, by constructing an amino acid substitution matrix for each motif, we assessed the potential pathogenic impact of every possible missense variant at each position within these motif classes.

## Materials and Methods

### ELM motif classes and regular expressions

Motif classes from the ELM resource were selected based on the following criteria: they had to have a fixed length, at least one experimentally validated instance in humans (true positive), and a minimum of five pathogenic or benign variants. Classes associated with post-translational modifications were excluded. The selected classes were subdivided into those with a crystallographic structure available in the Protein Data Bank (PDB), and those without. The former set was used in the AF2 motif-domain complex structure prediction validation procedure. The defining regular expressions for these classes were written following the nomenclature specified in the ELM database.

### SLiMs identification in the Human proteome

The whole human proteome was defined as the 20,435 canonical protein sequences with Reviewed status (i.e., Swiss-Prot) in the UniProt database (as of September 2024) [20]. SLiM regular expression matches were identified using an in-house-developed Python script that scanned the entire sequences, allowing multiple hits per sequence. Matches were defined as occurrences of the SLiM regular expression within the analyzed protein, characterized by the specific sequence, its position, and its length. The complete set of all such matches identified across the human proteome for a given motif class is called the “Initial Set”.

### Identification of naturally occurring variants within each motif

Clinically relevant missense variants involved in SLiM analysis were extracted from the ClinVar (FullRelease, September 2024) and gnomAD (v4.0 - Exomes, November 2023) databases. Only variants classified in ClinVar as Pathogenic, Pathogenic/Likely Pathogenic, Likely Pathogenic, Benign, Benign/Likely Benign, or Likely Benign were considered. Clinical significance and the number of corresponding supporting submissions were recorded for these variants. Benign missense variants exceeding a predefined allele frequency threshold were retrieved from gnomAD. For genes with at least 10 pathogenic variants, the allele frequency threshold was determined based on the most frequent pathogenic variant reported in ClinVar. For genes with fewer than 10 pathogenic variants, thresholds based on autosomal recessive (AR) and autosomal dominant (AD) inheritance models were applied (10^-4.1 and 10^-4.28, respectively), informed by the respective allele frequency distributions of pathogenic and benign variants in ClinVar. Finally, for genes not yet associated with any disease, a fixed threshold of 10^-4.1 was applied. The threshold values varied from 0.7637 to 0.000015, and are those commonly used in our group for variant annotation, following ACMG criteria [21].

### Structure data set

The crystallographic structures associated with motif instances of ELM classes were retrieved from the PDB. For ELM classes without associated crystallographic structures, 10 predictive models were generated using AlphaFold2, with the motif and domain sequences of experimentally validated human instances (i.e true positives) from the ELM database as input. These models were computed using two different settings for the number of recycles: 24 (five models per class) and 72 (five models per class).

### Stability Free-Energy Change calculations

The stability Gibbs free-energy change (△△G) for each missense variant was calculated using FoldX software for each structure of the corresponding motif-domain complex (derived from PDB or AF2). The calculations were performed using the PositionScan command. The resulting △△G values for each motif class are presented as structural stability free-energy substitution matrices. When multiple structures were available for a given complex, the matrix values represent the average △△G across all structures.

### Clinical Significance, Population Allele Frequency, and Free-Energy Change Conversion into Confidence Scores

The clinical significance of variants, as reported in ClinVar, was converted into confidence scores, where each review status star contributed 1 point, and each supporting submission added 0.1 points. Population allele frequencies from gnomAD and FoldX stability free-energy change values were also transformed into confidence scores using the respective equations described in Figure 2 SI.

### Conservation Score calculation

Multiple sequence alignments (MSAs) were constructed using sequences from the UniRef50 cluster associated with each match-containing protein. The alignments were performed using MAFFT software and subsequently used to calculate the motif conservation score, determined through the Jensen-Shannon Divergence algorithm [22,23].

### Secondary Structure prediction

Protein secondary structure prediction was performed using the Scratch software, with the full sequence provided as input. The output consisted of a per-residue classification into helix, strand, or other [24].

### Residues Exposure calculation

The solvent accessible surface area (SASA) was calculated using the FreeSASA library, and applied to the protein structure predictions provided by AlphaFold2 (AF2) for the SwissProt human proteome available on its website [25].

### Gene Ontology terms

Gene Ontology (GO) terms were retrieved from The Gene Ontology Resource [26].

### Protein structure and matrices visualization

Protein structure visualizations and substitution matrices were generated using Visual Molecular Dynamics (VMD) software and the Seaborn library [27,28].

### SLiMs analysis and filtering pipeline

A flowchart of the pipeline is provided in the Supplementary Information. The Initial SLiM set was filtered to retain only matches containing at least one variant from ClinVar or gnomAD, forming Set 0 (S0). Additionally, motifs experimentally confirmed in the ELM database were classified as Positive Set 0 (P0), representing the true positives. These motifs were analyzed for secondary structure, solvent exposure, conservation and associated GO terms. Clinical significance matrices were constructed using the variant data for each protein in P0, while structural stability matrices were generated using FoldX applied to PDB or AF2 structures.

Once the matrices were built, motifs were filtered in a three-step iterative cycle:

1. **Clinical Significance Cycle**: for all motifs in S0 their variants were classified as pathogenic or benign using clinical significance from ClinVar or gnomAD allele frequency thresholds. Motifs showing no contradictions against the true positive set (i.e. passing both the variant matrix and features evaluations) result in Positive Set 1 (P1). Motifs with discordant variant classifications were assigned to Negative Set 1 (N1), while unresolved cases were allocated to Remaining Set 1 (R1). Variants within motifs in P1 were used to refine ClinVar and gnomAD matrices. The cycle iterated until no new positive motifs could be identified.
2. **Structural analysis Cycle**: motifs in R1 were evaluated by comparing their variants against the FoldX stability matrix obtained from the corresponding motif-receptor complex structures. Tolerated SAS are those whose ΔΔG is below predefined thresholds of 2.1 kcal/mol for PDB structures and 1.6 kcal/mol for AF2 models (both calibrations are shown in Supplementary Information, Fig 3 SI and Fig 4 SI). Motifs whose variants do not contradict the resulting SAS tolerance matrix were assigned to Positive Set 2 (P2). Discordant and unresolved cases were transferred to Negative Set 2 (N2) and Remaining Set 2 (R2). As for the clinical significance cycle, motifs in P2 were used to further refine the matrices. In this cycle, only the well-defined residue positions present in the regular expression are analyzed. Flexible positions (i.e. x or ^P) in the regular expressions are analyzed in the last cycle.
3. **Flexible position Cycle**: motifs in R2 were finally filtered analyzing the SAS (in)tolerance as evaluated using FoldX ΔΔG values in the regular expression flexible positions. The methodology mirrored that of the previous cycles, ultimately resulting in the final sets: Positive Set 3 (P3), Negative Set 3 (N3), and Remaining Set 3 (R3).

Motifs in P3 were designated as functional with high confidence, while those in N3 were classified as non-functional with equal confidence.

## Results

### 1) Alpha fold provides high-quality SLiM-receptor complex structures

While the overall quality of AF2 models has been extensively tested, igniting a plethora of studies based on those structures, it remains essential to evaluate whether the generated predictions are suitable for the particular purposes for which they are intended [29–31]. In line with this reasoning, we first used AF2 to replicate the core motif-receptor structures of 17 ELM motif classes (the same group analyzed in our previous work, and some additional cases), each of which had at least one crystallographic structure deposited in the PDB using only their sequence information as input. Kery to our approach is the accurate positioning of the SLiM within its corresponding binding site (Figure 1A, left panel). Analysis of AF2 models reveals that, while AF2 reliably predicts the overall receptor structure, improper SLiM positioning is observed in certain cases (red SLiM in the model shown in Figure 1A). This mispositioning often leads to AF2 failing to identify the correct motif binding site.

**Figure 1.**
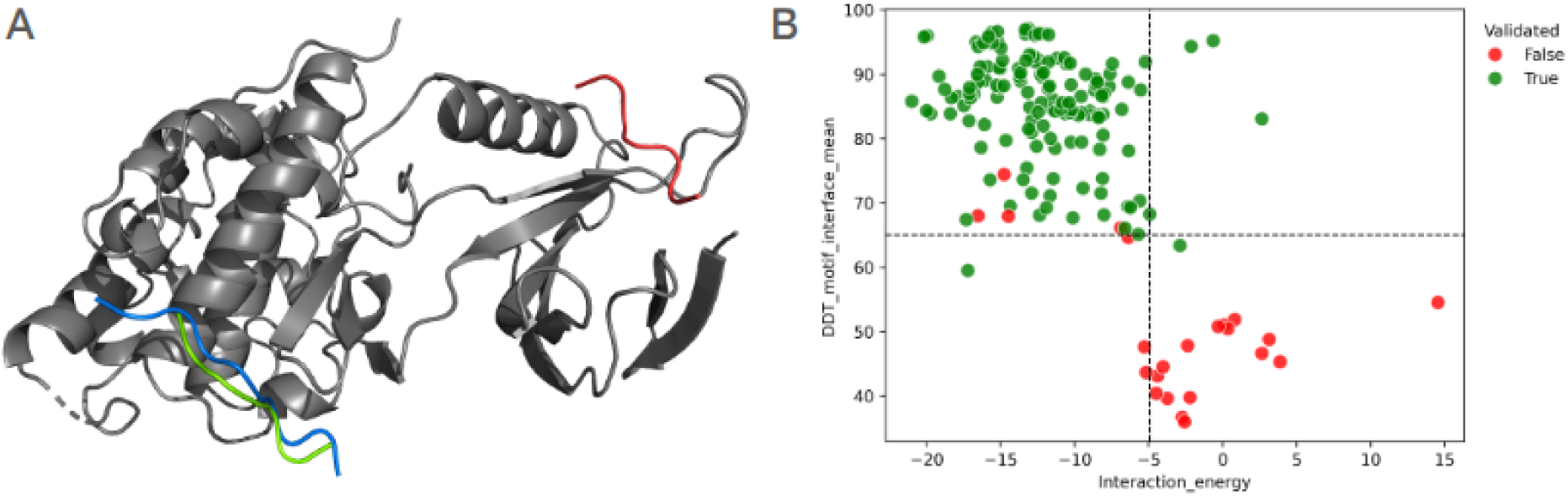
(A) Crystallographic structure (pdbid 1UKH), depicting the interaction between the DOC_MAPK_JIP1_4 motif (in blue) of mouse JIP1 (UniProtID: Q9WVI9) and the Pkinase domain (in grey) of human MAPK8 (UniProtID: P45983). Two AF2 predictions for the motif are shown: a correct prediction (in green), and an incorrect prediction (in red). (B) Scatter plot showing the relationship between the mean pLDDT of motif residues (y-axis) and the interaction energy in kCal/mol (x-axis). Each dot represents an AF2 prediction: green dots correspond to predictions classified as reproducible based on manual inspection and their agreement with the respective crystallographic structure in the PDB, while red dots represent incorrect predictions. Dashed lines indicate threshold values selected to distinguish between correct and inaccurate AF2 predictions.

To address this issue and avoid using motif-receptor complexes with incorrectly positioned motifs, we analyzed several parameters of the AF2 models to assess their ability to discriminate between correct and incorrect SLiM positions. As shown in Figure 1B (right panel), correct models exhibit a higher mean pLDDT for AF2-predicted SLiM residues and a lower interaction energy between the receptor and the SLiM. Based on these observations, we defined threshold values of 65 for the mean pLDDT of the motif residues, and −5 kcal/mol for the interaction energy. The former reflects AF2’s self-assessed confidence in positioning the motif residues at that location relative to their domain binding partner, while the latter accounts for the thermodynamics contributions facilitating the complex formation. In other words, once several AF2 models are built for a given SLiM-receptor complex (we generated 10 models per complex), selecting the one with the highest mean pLDDT for motif residues and the lowest binding energy provides the optimal selection criterion, provided it meets the defined thresholds. Indeed, complex structures with properly positioned SLiMs could be selected using only these two parameters in all tested cases.

At the heart of MotSASi are the structural SAS tolerance matrices. We thus compared the corresponding matrices obtained from the crystallographic structures deposited on PDB with those derived from the AF2 models that passed the previous quality checks. Visual inspection of the two matrices (shown in Supplementary Information, Figure 5 SI) clearly reveals that they are highly similar. To quantify this similarity, we used two metrics. First, we computed the Pearson correlation coefficient between the ΔΔG values obtained with either matrix, yielding a value of 0.65 +/- 0.22. Secondly, we classified each SAS as tolerable or non-tolerable for each matrix, defining those derived from the crystal structure as the reference (or true) cases and those from AF2 as the predicted cases. The resulting statistical performance values (e.g. an accuracy of 0.79) demonstrate that AF2 models are sufficiently accurate to be incorporated into the MotSASi pipeline.

### 2) SLiM natural variant analysis supports AlphaFold2 models

Having shown that AF2 models yield accurate SLiM-receptor structures and their resulting SAS tolerance matrices, we further analyzed whether examining known SLiM variants could aid in selecting the correct AF2 complex models. Comparing the FoldX stability matrix with the observed variant (from ClinVar and gnomAD) pathogenicity matrix provides an independent and stringent test of the AF2 model. Figure 2 presents a visual example of the matrix comparisons for correct, and incorrect models.

**Figure 2.**
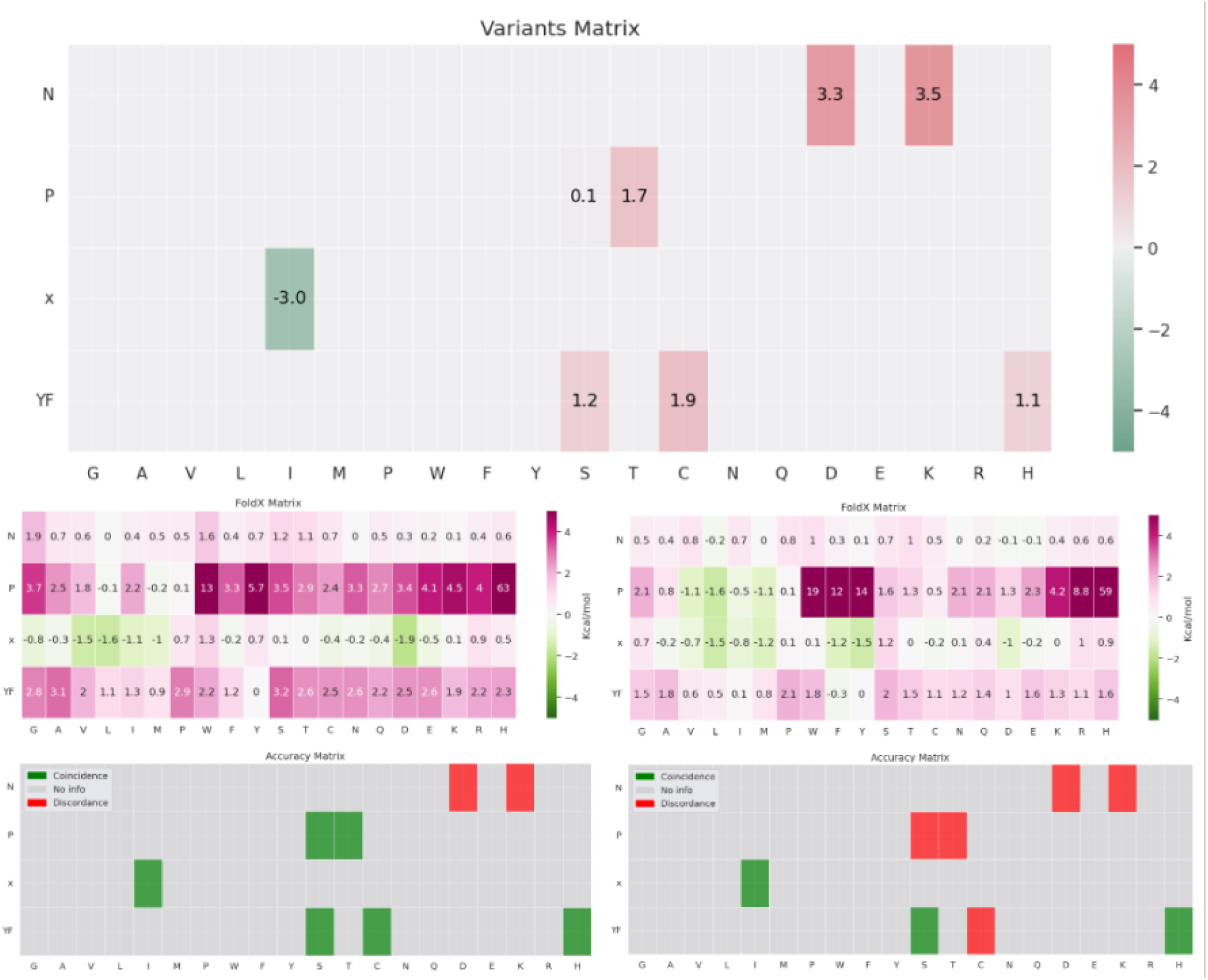
Comparison of the observed variant pathogenicity matrix (upper panel) with FoldX stability matrices for the correct (left panel) and incorrect AF2 models (right panel).

Interestingly, although the structures and positioning of the SLiM relative to the receptor differ completely between the two models, the matrices still display some similarity. This observation arises from the inherent properties of residues and their general impact on protein structure (e.g. substituting a rigid Proline at row 2, as shown in the example of Figure 2, is typically more destabilizing than mutating the degenerate residue in row 3). It further underscores the importance of carefully comparing the FoldX stability matrix with the observed variant pathogenicity matrix. This comparison is illustrated in the lower panel using a binary color scheme: green cells indicate agreement between matrix predictions, while red cells indicate disagreement. The correct model achieves 6 matches out of 8 tested variants, whereas the incorrect model achieves only 3. Similar trends are observed for other motifs. Unfortunately, as previously discussed, observed variant pathogenicity matrices tend to be scarce (i.e. they include data for only a limited subset of possible variants), making them unsuitable for individual assessments. Instead, we performed a broader evaluation by aggregating all observed variants to calculate a global accuracy metric. Correct models achieved an accuracy of 74.72% while incorrect models achieved 63.16%.

Summarizying both analysis, it is clear that taken together the model pLDDT, the SLiM-receptor interaction energy and the comparison of the SAS and clinical significant matrices, allow selecting the best AF2 model.

### 3) AlphaFold2 allows increased high-confidence SLiMs prediction

One of the main limitations of the previous version of MotSASi was the scarcity of motif-domain structures available in the PDB. This limitation constrained both the computation of FoldX stability matrices, and the identification of new high-confidence candidates. The previous version of MotSASi included 20 different SLiM-receptor complexes, encompassing 4,389 analyzed motifs hosted by 3,085 unique proteins. By updating the databases and incorporating AF2-generated models of motif-receptor complexes, we have now analyzed 51 distinct ELM motif classes—27 with crystallographic structures deposited in the PDB and 24 based on AF2-generated models—resulting in a total of 8,731 analyzed motifs corresponding to 5,027 unique proteins (Table 1). Considering that the current human proteome in SwissProt comprises 20,435 entries, our predictions cover approximately 25% of the human proteome. Furthermore, among the 4,866 gene-encoded proteins associated with Mendelian diseases reported in OMIM, 1,694 contain at least one high-confidence motif, increasing the coverage to 35% of disease-associated proteins. This information is summarized in Table 1.

**Table 1.** Summary of the analyzed ELM motif classes, instances, and associated human proteins.

|  | MotSASi | PDB | AF2 | OMIM |
| --- | --- | --- | --- | --- |
| # ELM classes | 51 | 27 | 24 | 42 |
| # ELM instances | 405 | 305 | 100 | 199 |
| # RE proteome hits | 377,170 | 165,481 | 211,689 | 124,171 |
| # No variants motif instances<br>(% of total) | 315,957<br>(84%) | 137,444<br>(83%) | 178,513<br>(84%) | 98,554<br>(79%) |
| # Negative motif instances<br>(% of total) | 52,482<br>(14%) | 22,218<br>(13%) | 30,264<br>(14%) | 22,047<br>(18%) |
| # Positive motif instances<br>(% of total) | 8,731<br>(2%) | 5,819<br>(4%) | 2,912<br>(1%) | 3,570<br>(3%) |
| # Proteins | 5,027 | 3,896 | 2,360 | 1,694 |
| Enrichment against ELM inst. | X 21.56 | X 19.08 | X 29.12 | X 17.94 |

The MotSASi pipeline presented herein, with the key inclusion of AF2 models, expands the coverage of human ELM instances nearly 22-fold, with enrichment levels ranging from 19-fold for the PDB group to 29-fold for the AF2 group. Encouragingly, the pipeline discards approximately 14% of regular expression matches as potential false positives, deeming them unlikely to represent functional SLiMs. Nonetheless, significant limitations persist, as approximately 84% (315,957) of the SLiM regular expression matches processed by MotSASi lack known variants and, as such, cannot yet be fully analyzed. Despite these challenges, the increased coverage of high-confidence potential SLiMs and the identification and elimination of false positives are expected to improve further as more variants are discovered and classified.

#### Helical calmodulin-binding motif

As an example of a newly analyzed ELM class using AF2, we present the LIG_CaM_NSCaTE_8 motif (ELM RegEx: W[^P][^P][^P][IL][^P][AGS][AT]). Interestingly, two experimentally validated human instances of this motif were previously reported in the ELM database, located in the alpha subunit of L-type voltage-gated calcium channels (VGCCs). One instance is found in the N-terminal cytoplasmic domain of Cav1.3 (HGNC gene name: *CACNA1D*, UniProtID: Q01668) and is known as the N-terminal spatial Ca²⁺-transforming element (NSCaTE) motif. This motif has been extensively studied and shown to play a key role in accelerating channel closure when the N-lobe of Ca²⁺-CaM binds to the NSCaTE motif, a regulatory mechanism called calcium-dependent inactivation [32]. Loss-of-function variants in *CACNA1D* are responsible for a Mendelian disorder known as SANDD (SinoAtrial Node Dysfunction and Deafness, MIM Number: 614896), which follows an autosomal recessive inheritance pattern [33]. Notably, *CACNA1D* is expressed in neuronal cells and cardiac myocytes.

Through our iterative analysis, we were able to discard 74 motifs and identify 7 new high-confidence potential candidates for this SLiM in the human proteome. For instance, Ca²⁺-CaM, a ubiquitously expressed protein critical for various cellular functions, interacts with numerous targets. These targets generally consist of sequences of 15–30 amino acids with an intrinsic propensity to form an alpha helix. Our AF2 model and the accompanying amino acid substitution matrix provide a deeper understanding of the motif-domain binding event. As shown in Figure 3, Trp53 occupies a central position within the hydrophobic pocket formed by the EF-hand of CaM. Regarding the substitution matrix, we observe that Trp in the first position, Ile or Leu in the fifth position, and small hydrophobic amino acids in the eighth position are highly constrained. These positions exhibit limited allowance for alternative amino acids. In contrast, the fourth position, previously defined as highly degenerate by ELM, appears less permissive, favoring only small hydrophobic residues. Based on our SAS matrix analysis (Figure 6 SI), we propose the following refined regular expression for the NSCaTE motif: W..[APC][VLIM][^P][^P][GAVMSTC].

**Figure 3.**
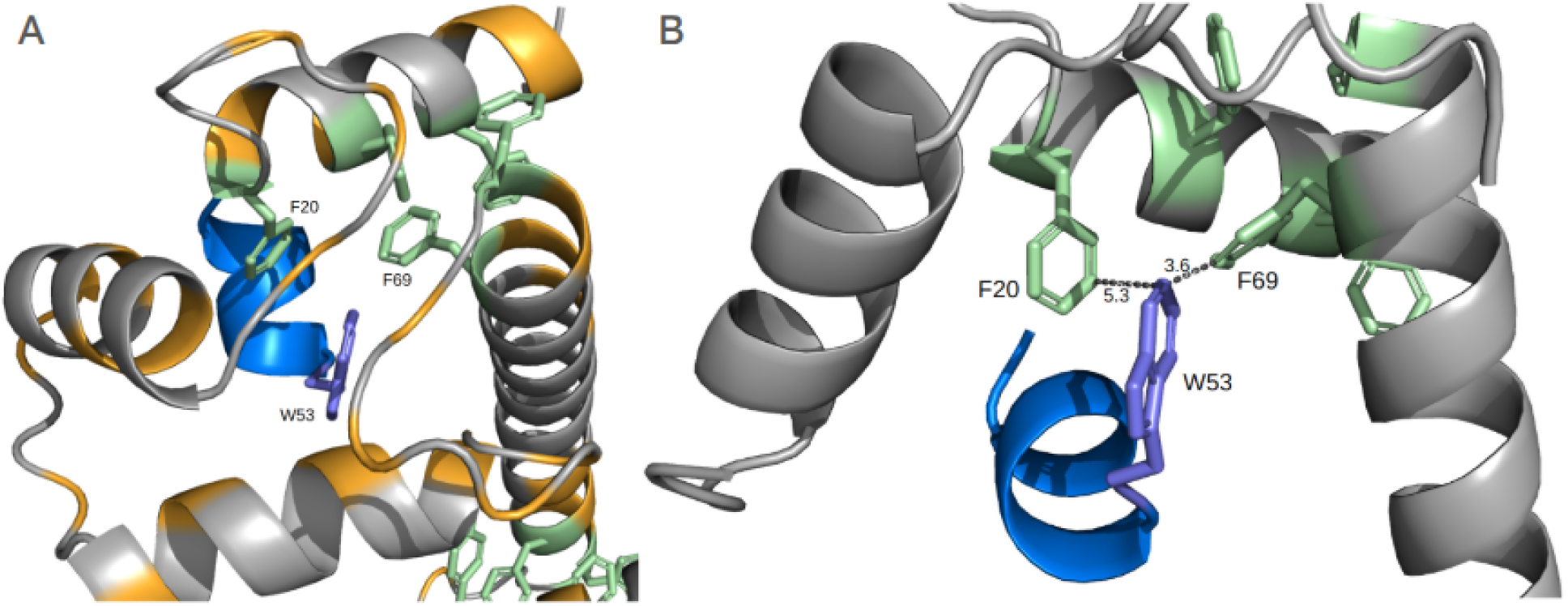
(A) Trp53 (shown in purple) of Cav1.3 (UniProt ID: Q01668) occupies a central position within the hydrophobic pocket formed by the EF-hand of CaM (UniProt ID: P0DP25). The motif chain is shown in blue, the domain chain in grey, aromatic residues of the domain in green, and aliphatic residues of the domain in orange. (B) Trp53 viewed from another angle, with distances to neighboring aromatic residues depicted (measured in Å).

### 4) SLiM-receptor complex models improve SLiM variant pathogenicity prediction

MotSASi, as an algorithm, leverages clinical significance and population allele frequency data of variants reported in relevant databases. Nevertheless, integrating crystallographic structures to facilitate thermodynamic predictions of variant deleteriousness remains critical. In this context, we sought to compare MotSASi with another method that relies on structural information to train its algorithm. Given its recent success, we selected AlphaMissense (AM) as the candidate. To ensure a fair comparison, we randomly set aside a subset comprising 20% of ClinVar pathogenic and gnomAD benign variants found in ELM instances (true positives) and/or newly identified high-confidence motifs, to serve as an independent test set. The final test set consisted of 2,628 variants, including 2,449 gnomAD benign and 179 ClinVar pathogenic variants. MotSASi outperformed AlphaMissense across all three evaluated metrics: accuracy (98% vs. 76%), F1-score (0.99 vs. 0.85), and Matthews Correlation Coefficient (MCC) (0.83 vs. 0.33). It is clear that while AM is an excellent general-purpose variant pathogenicity prediction tool, the detailed analysis of underlying structures in a biological context, as performed by MotSASi, significantly improves overall prediction accuracy.

**Table 2.** Confusion matrices comparing the performance of MotSASi and AlphaMissense on a validation set of variants.

|  | Pathogenic<br>MotSASi verdict | Benign<br>MotSASi verdict |  | Pathogenic<br>AM verdict | Benign<br>AM verdict |
| --- | --- | --- | --- | --- | --- |
| ClinVar<br>Pathogenic | 130 | 49 | ClinVar<br>Pathogenic | 152 | 27 |
| gnomAD<br>Benign | 5 | 2444 | gnomAD<br>Benign | 607 | 1842 |

Analysis of the confusion matrix shows that the key to the observed improvement in MotSASi lies in its ability to avoid tagging known benign variants as pathogenic. The naturally imbalanced dataset, with many more benign variants as often observed in real-world examples, results in AM misclassifying 607 benign variants as pathogenic, whereas MotSASi shows only 5 cases of these potential false positives. As expected, for known pathogenic variants, while MotSASi fails to detect 49, AM only fails in 27 cases. Generally speaking, it appears that MotSASi tends to be more conservative than AM concerning pathogenic classification, which aligns with current ClinGen recommendations.

To gain deeper insight into the comparison, we examined specific cases. A well-studied example is the LIG_PTB_Apo_2 motif in LDLR, which is crucial in its internalization upon ligand binding. The motif follows the regular expression NP.[YF], and the MotSASi-defined SAS tolerance matrix allows refining it to N[PA][^GWFYKH][YFWML]. Several variants affecting this motif have been previously reported in ClinVar and gnomAD. At the first position, Asn825Asp and Asn825Lys variants have been reported as pathogenic. The substitution to lysine is correctly predicted by AM as Likely pathogenic, whereas aspartate is incorrectly classified as Likely benign. At the second motif position, Pro826Ser and Pro826Thr have been reported as pathogenic, and again the first variant is correctly predicted as Likely pathogenic by AM, while the latter is classified as Uncertain significance. The third motif position lacks clinically significant variants; however, a Leu827Ile variant with a high allele frequency is reported in gnomAD. It is correctly classified as Likely benign by AM. At the fourth and final motif position, three missense variants (Tyr828Ser, Tyr828Cys, and Tyr828His) have been reported. All three are classified as Uncertain by AM. In contrast, MotSASi correctly predicts all these variants by leveraging the information of the SAS tolerance matrix.

This misclassification could stem from the fact that AM bases its predictions on individual protein structure assessments without explicitly considering protein-protein binding events. Inspection of the 1NTV crystallographic structure deposited in the PDB, which represents the corresponding motif-domain interaction, can help us better understand this trend. For the misclassified Asn825Asp variant, we observe that the mentioned Asn is involved in a double hydrogen bond. While its oxygen donates an electron pair to the backbone nitrogen of the Val at the third position, it simultaneously acts as a hydrogen donor through its amide nitrogen with a backbone oxygen from the domain chain. Additionally, there is a Phe residue in close proximity, which most likely is stabilized in the neutral environment. Therefore, it seems highly unlikely that a negatively charged residue, such as Asp, could be tolerated, leading to the observed pathogenic variant. Regarding the second motif position, it appears that the proline fulfills a specific stereochemical role, due to its particular backbone conformation, enabling the association between the first and third positions while occupying minimal space. It seems reasonable to assume that only Pro would be accepted, even though MotSASi’s energetic considerations suggest that other small aliphatic residue might also be possible. At the third position, occupied by a Val, we find it in close proximity to hydrophobic residues. This suggests that not all residues would be allowed, but rather only aliphatic ones of a specific size. AM fails to recognize this and cannot predict the pathogenicity of either Thr or Ser substitutions. Finally, at the fourth position, the Tyr residue appears to occupy a pocket adapted for an aromatic residue, adding an hydrogen bond between the Tyr hydroxyl group and the carbonyl oxygen in the domain backbone. As reflected by MotSASI SAS tolerance matrix it seems reasonable to conclude that mutations to non-aromatic residues would likely disrupt this interaction.

## Discussion

In this work, we present our progress in studying SLiMs in human proteins, leveraging increasingly enriched variant databases, and the recently developed Artificial intelligence (AI) algorithms for protein structure prediction. Our focus, oriented towards the broad spectrum of Mendelian disorders, specifically addresses missense variants located in SLiMs. These functional elements, which play key roles in numerous physiological processes, are vulnerable to disruption by genetic alterations. When these changes occur in genes associated with a given disease, they can lead to the development of the underlying pathology, making them clinically relevant.

Our pipeline significantly expanded the repertoire of known functionally relevant SLiMs, achieving an almost 22-fold increase in the number of analyzed motifs in the human proteome. While this progress is still far from capturing the entirety of potential functional SLiMs—predicted to be around one million—it represents an important step in providing experimental scientists with hypotheses and advancing our understanding of protein-protein interactions. As highlighted in our previous work, a key strength of the MotSASi method lies in its ability to integrate reported missense variants and three-dimensional protein structural data to generate more accurate predictions. Conversely, it is crucial to emphasize the value of filtering out non-functional motifs, as this prevents researchers from expending unnecessary effort on dubious Variants of Uncertain Significance (VUS). Given the clinical focus of MotSASi, we restricted our analysis to motif matches associated with clinically interpretable variants. Our method showed comparable performance when using both PDB and AF2 structural inputs.

Generating AF2 predictions introduces another layer of complexity. Although ELM provides curated data, gaps often require manual intervention. For example, missing interaction pairs between motifs and binding domains, outdated transcript references, and alternative (no-Pfam) domain annotations complicate the preparation of FASTA files for AF2 predictions. In some cases, generating an AF2 prediction with the desired minimum quality levels of motif-domain binding energy and pLDDT confidence values for motif residues is impossible. Biological challenges also arise, such as motifs requiring post-translational modifications (e.g., phosphorylation), which AF2 cannot currently model. Attempts to use phosphomimetic residues to approximate phosphorylated states in PDB structures yielded poor results when phosphates were critical for binding. Similarly, motifs involving non-standard stoichiometries, such as interactions with dimeric domains, posed difficulties. Nevertheless, we successfully addressed these cases by modeling sequences with domain repeats, demonstrating AF2’s utility when provided with accurate input.

The high-confidence SLiM candidates and missense substitution matrices provided by MotSASi contribute to the clinical genomics community with tools to refine ACMG guidelines for variant interpretation, particularly the PM1 label [34]. PM1 label is assigned for those variants “located in a mutational hot spot and/or critical and well-established functional domain without benign variation”. This label has not been reviewed by ClinGen experts in recent years, which is a gap of concern for our group. Traditionally, PM1 candidate residues or regions were assigned based on known enzyme active sites, recurrent mutational “hot spots” or other protein residues/regions with a well-established functionality usually including single mutagenesis experiments. SLiMs are good candidates as small functional domains, i.e. PM1 candidates. However, even though it may be tempting to consider the whole motif for PM1, our results clearly show that further refinement is needed, and possible as shown by MotSASi. Our assignment of high-confidence motifs combined with the final SAS tolerance matrices allows us for each human protein with high confidence SLiM to determine which residues (and which SAS) are non-tolerant, and thus their underlying variants should be labeled with PM1. In this context, and to help clinical genomic analysts implement this labeling criteria, we provide a file (as part of the SI) with all predicted variant effects in high-confidence SLiM candidates and known instances cataloged in the ELM database. In other words, in a clinical setting, we propose assigning PM1 to variants located within high-confidence functional motifs and a high non-tolerant confidence score value in the final SAS matrix.

Finally, given our strong emphasis on the structural component of motif-domain interactions, we compared MotSASi with AM. Previous studies evaluating AM indicate that the algorithm performs less accurately in disordered regions, as defined by AF2 using residue confidence scores below a specific threshold [35]. This reduced performance is likely due to the enrichment of benign variation in disordered proteome regions, which paradoxically also harbor functional sequences such as SLiMs, where pathogenic variants can disrupt motif-domain interactions. Although MotSASi demonstrated markedly higher accuracy than AM (98% vs. 76%), this result may be misleading due to the high enrichment of benign variants in the test set. The literature has extensively discussed this issue, particularly regarding the typical imbalance between positive and negative instances in clinical genomics datasets [36]. In such cases, alternative metrics are recommended, such as the Matthews Correlation Coefficient (MCC), which rewards methods that correctly predict both positive and negative instances. Using this metric, MotSASi also outperformed AM (0.83 vs. 0.33). Analysis of the confusion matrices generated by each method revealed that, while AM provides more accurate predictions for pathogenic variants in the test set, it also produces many false positives. Beyond the “data problem,” it is important to emphasize that in clinical genomics, false negatives are generally preferable to false positives. Extrapolated to molecular diagnosis, it is more prudent to withhold reporting a variant until stronger evidence is available, rather than report prematurely and risk subsequent retraction.

In summary, after the first MotSASi publication, we predicted that advancements in massive sequencing technologies, their integration into clinical practice, increasing experimental validation of predicted SLiMs, and new ligand-domain crystal structures would collectively enhance our understanding of amino acid substitution effects within motifs. This, in turn, would lead to more precise and accurate predictions of biologically relevant SLiMs. Remarkably, most of these developments have materialized in just three years—a surprisingly short time frame. Envisioning the future possibilities and direction of cutting-edge SLiM analysis remains challenging. Will refined AF2-like predictions for individual motif-domain pairs take center stage? Could multiplex assays of variant effects (MAVEs) be applied to SLiM studies, contributing directly to functional assay evidence for clinically relevant variants? Will large-scale functional assays identify most functional motifs across the human proteome? Until that time comes, significant efforts will need to be sustained, and MotSASi will play a pivotal role in striving to achieve the highest possible diagnostic yields in clinical genomics.

## Acknowledgements

To Mariano Martin, who contributed significantly to the development of MotSASi in its early stages, both in its theoretical biological foundation and its implementation at the programming level.

