## Supplementary Information for "Integrating AlphaFold2 Models and Clinical Data to Improve the Assessment of Short Linear Motifs (SLiMs) and Their Variants’ Pathogenicity"

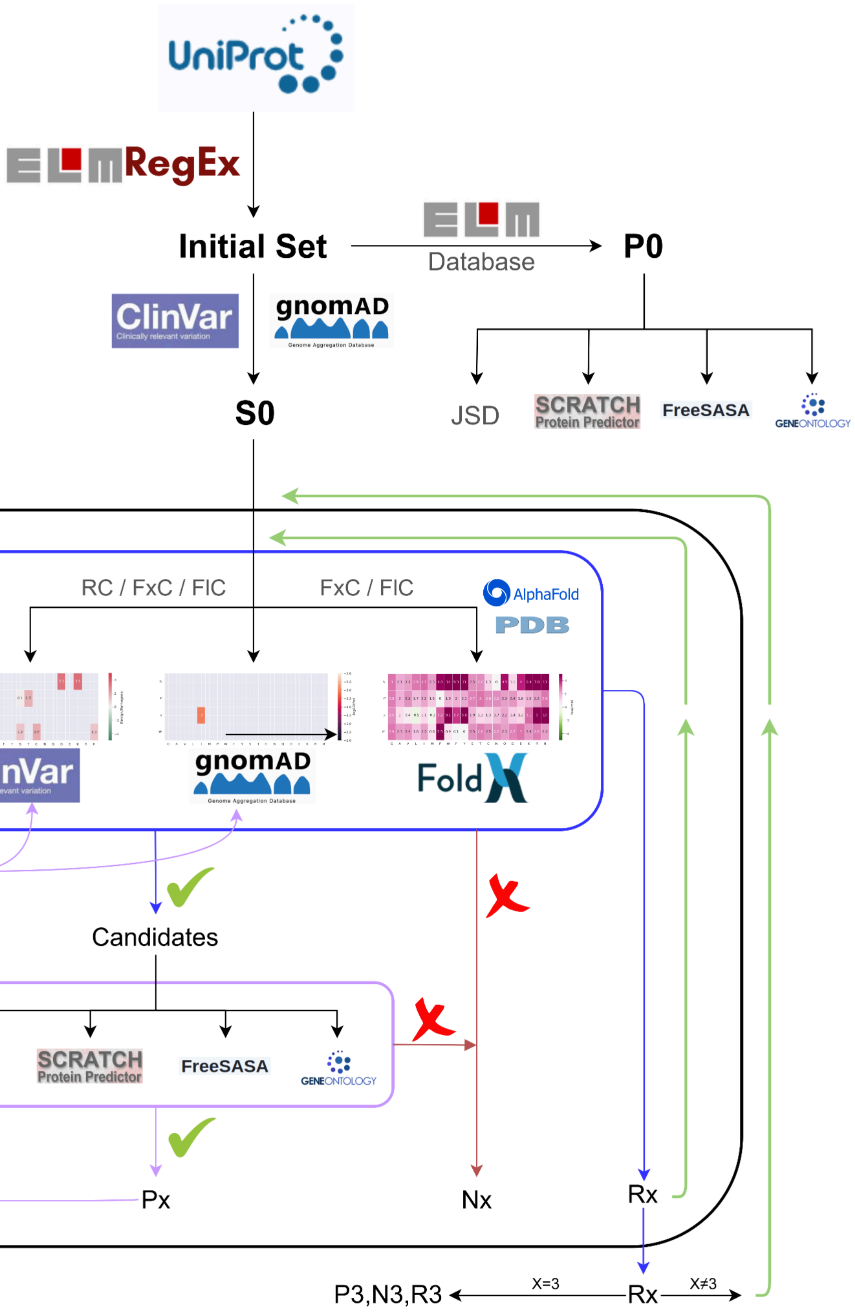


**Figure 1 SI:** Flowchart of the MotSASi pipeline, integrating crystallographic structures from the PDB and predictive models from AlphaFold2.

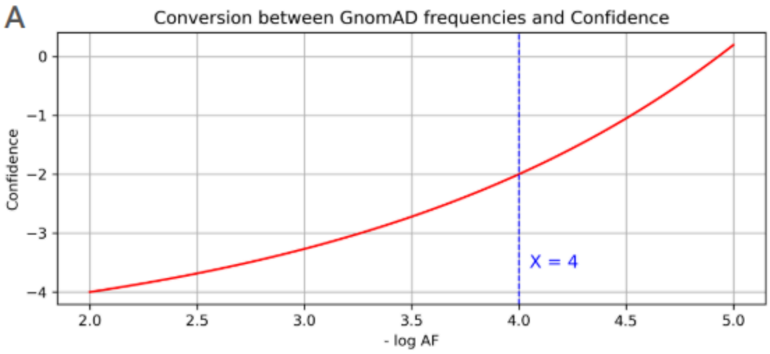


$$\text{Confidence Score} = \frac{1}{3} \cdot \left(\sqrt{3}\right)^{\log(\mathrm{AF})} - 5$$

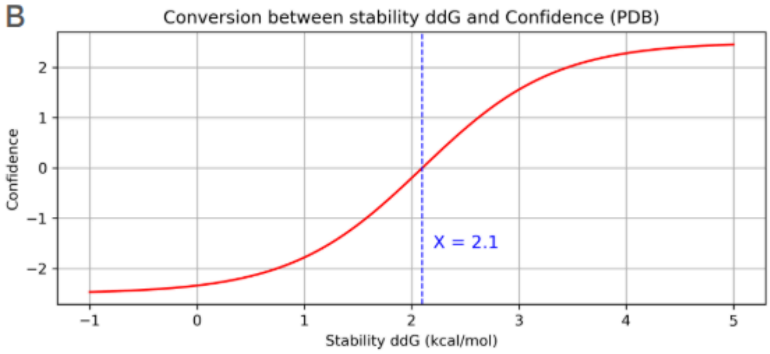


$$\text{Confidence Score} = \frac{5}{1 + e^{\ln\left(\left(\frac{2}{3}\right)^{4}\right)\cdot(x-2.1)}} - 2.5$$

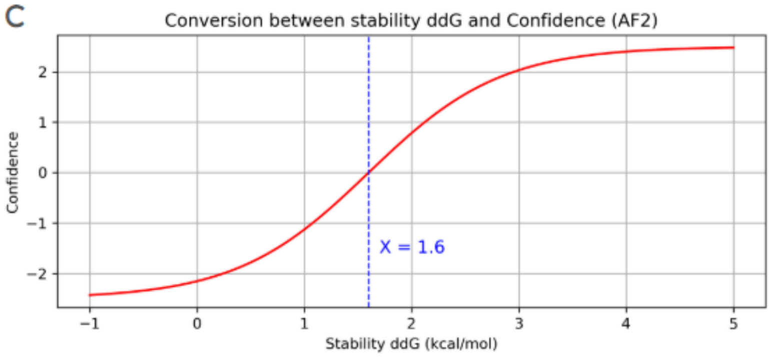


$$\text{Confidence Score} = \frac{5}{1 + e^{\ln\left(\left(\frac{2}{3}\right)^{4}\right)\cdot(x-1.6)}} - 2.5$$

**Figure 2 SI:** Plots of functions and their corresponding formulas for (A) gnomAD, (B) PDB-derived, and (C) AF2-derived Confidence Scores.

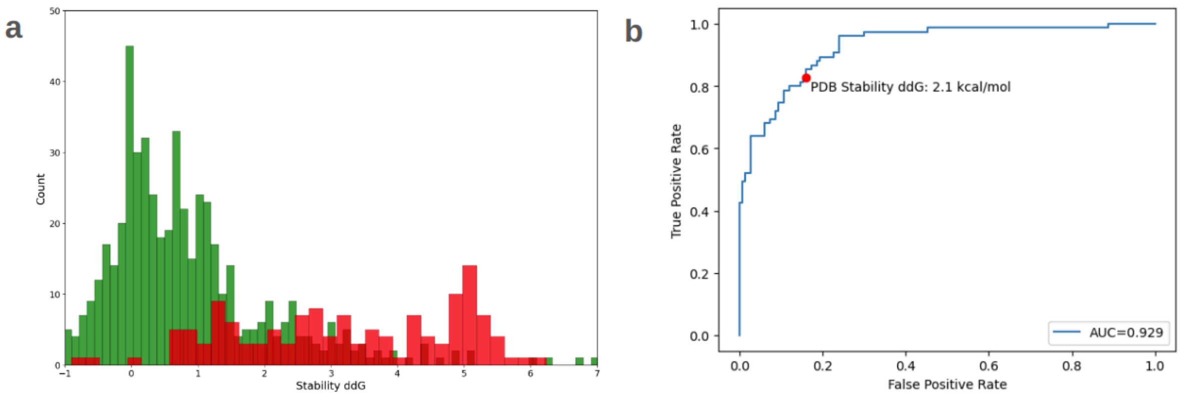


**Figure 3 SI:** (a) Histogram displaying the ClinVar and gnomAD variants used to define the ddG stability threshold. Pathogenic variants are indicated in red, while benign variants are shown in green. (b) ROC curve (AUC = 0.929) generated during the ddG threshold determination process for PDB crystallographic structures. The red dot marks the threshold value of 2.1 kcal/mol, corresponding to a sensitivity of 83% and a specificity of 84%.

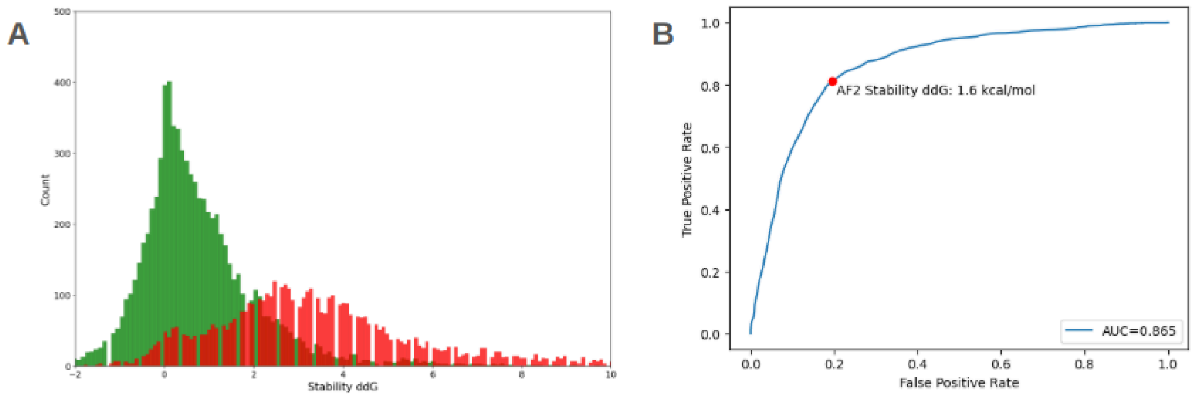


**Figure 4 SI:** (a) Histogram showing the ddG stability values calculated on crystallographic structures predicted by AF2 for variants previously classified as tolerated or non-tolerated in PDB-deposited structures. Non-tolerated variants are shown in red, while tolerated variants are shown in green. (b) ROC curve (AUC = 0.865) generated during the process of determining the ddG cutoff value for AF2.

The red dot marks the cutoff value of 1.6 kcal/mol, corresponding to a sensitivity of 81% and a specificity of 80%.

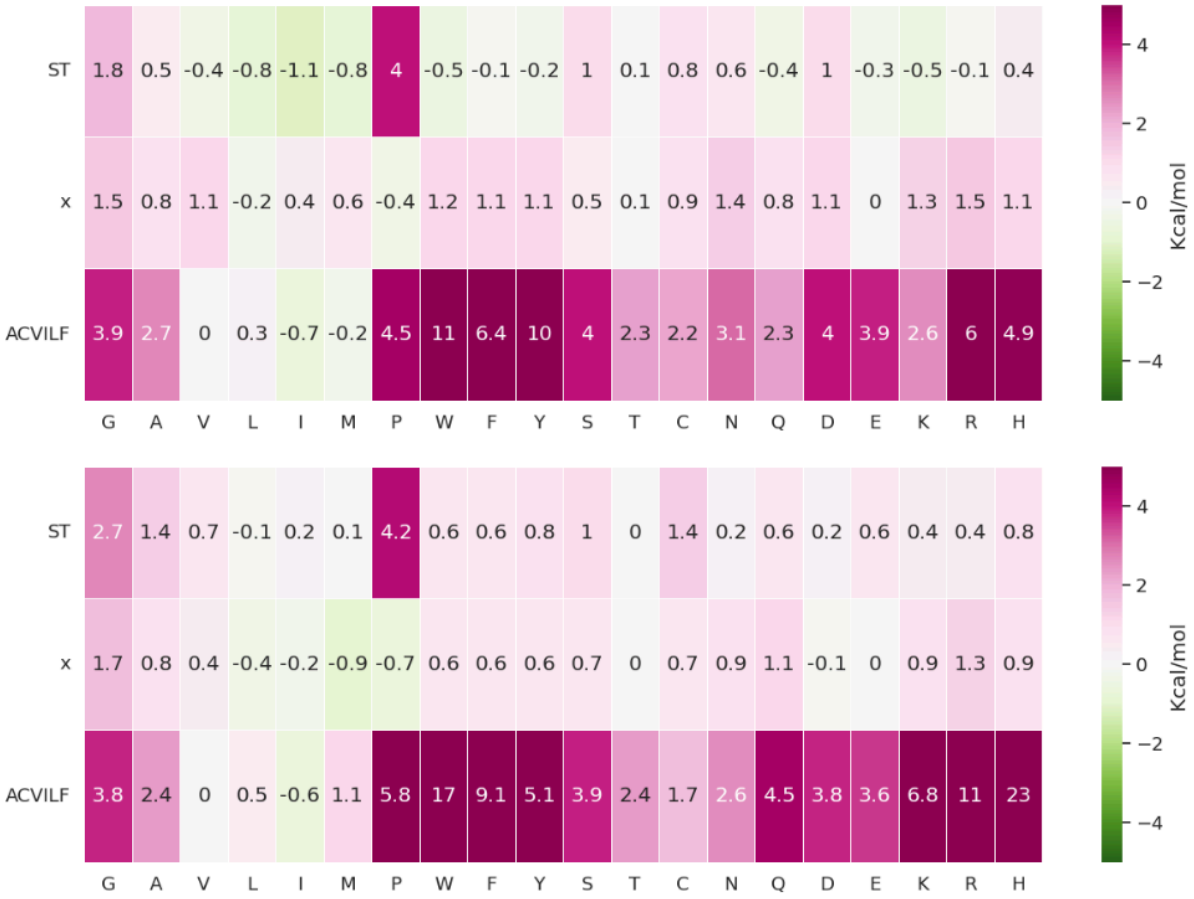


**Figure 5 SI:** Comparison of structural (stability ΔΔG) SAS matrices obtained using either the crystallographic structure deposited in the Protein Data Bank (PDB) (upper panel) or the SLiM-receptor structure modeled by AlphaFold2 (AF2) (lower panel).

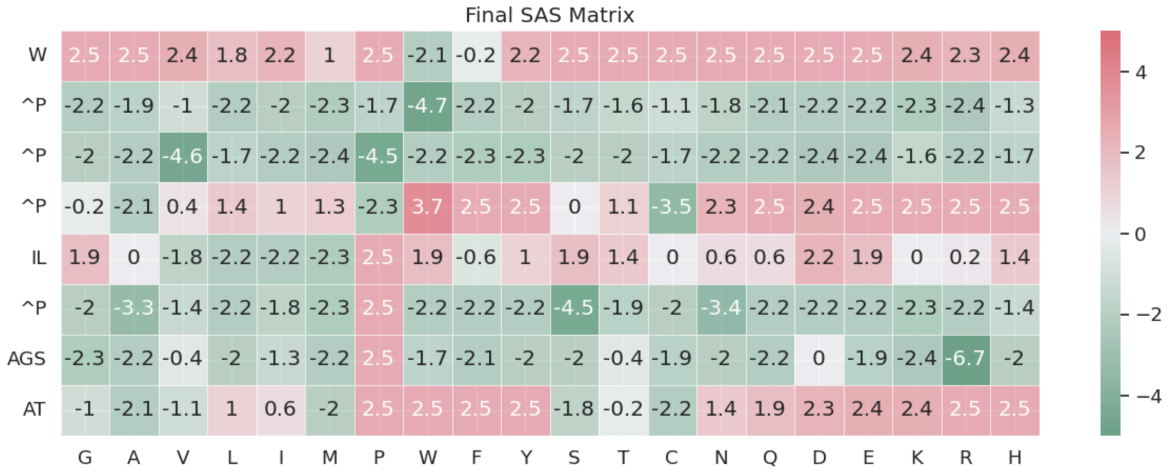


**Figure 6 SI:** Final SAS matrix with confidence scores in the cells for the NSCaTE motif.
